# Generating high quality reference genomes from field collected specimens by optimizing preservation

**DOI:** 10.1101/2023.03.10.532097

**Authors:** Jeremiah J Minich, Malia L Moore, Nicholas A Allsing, Emily R Murray, Loi Tran, Todd P Michael

## Abstract

Sample preservation challenges efforts to generate high-quality reference genomes or pangenomes for all 2 million+ plant and animal species. Here we show 95% ethanol protects against degradation for fish blood (22°C, ≤6 weeks) and plant tissue (4°C, ≤3 weeks). Using Nanopore, we assemble high quality reference genomes from three fish and two plant species (contig N50: 6.5-13.8Mb; BUSCO completeness: 94.4-99.2%; QV: 43.8 for *M. esculenta*).

## Main

Long-read sequencing, the 2022 Nature method of the year, is enabling biologists to digitally archive genomes from bacteria to complex eukaryotes to understand fundamental questions in evolution, bolster conservation strategies in the midst of mass extinctions, discover the biosynthetic machinery behind pharmaceutically relevant specialized metabolites, and improve food production through crop and livestock genomics.^1,2^ Large-scale initiatives such as The Earth Biogenome project and Darwin Tree of Life Project are employing this technology to generate reference genomes for all 1.8 million named eukaryotes on Earth including all 71,657 vertebrates by 2025.^3–5^

Sample collection, storage and preservation are critical steps to ensure high molecular weight (HMW) DNA acquisition required for long-read sequencing. Standard methods, which include snap freezing in liquid nitrogen or on dry ice followed by storage at -80°C, are often unrealistic in many Low and Middle Income Countries (LMIC), particularly at remote locations. Alternative storage methods (solvent, buffer, desiccation) that are compatible with short-read sequencing have shown theoretical promise in long-read sequencing applications through analysis of DNA fragment sizes and purity.^6^ However, these methods have yet to be benchmarked on a major long-read sequencing platform such as Oxford Nanopore Technologies (ONT) or Pacific Bioscience (PacBio), and have only been quality assessed at short storage times (hours) that are not realistic with travel times from remote regions.^6^ Improving the capacity to collect, store, and transport samples by using readily-available and cost-effective buffers is critical for the success of generating high quality reference genomes of all life on Earth along with supporting pangenome efforts to improve crop production.

Here we evaluate the impacts of storage solutions (95% EtOH and RNAlater), temperature (4°C and 22°C), and time (0 days, 4 hours, 2 days, 1 week, 3 weeks, and 6 weeks), on HMW DNA quality, ONT sequencing quality (LSK114, R 10.4.1) and finally assembly quality (Figure 1). We sampled fish (n=10 species, 100 samples) and plants (n=4 species, 36 samples) to establish the protocols across the tree of life (Supplementary Table 1). These complementary groups of organisms are realistically challenging to preserve in the field and represent different tissue types. We confirm the viability of solvent-preserved samples by producing high-quality *de novo* assemblies for five organisms at the furthest storage time point.

**Figure 1.**
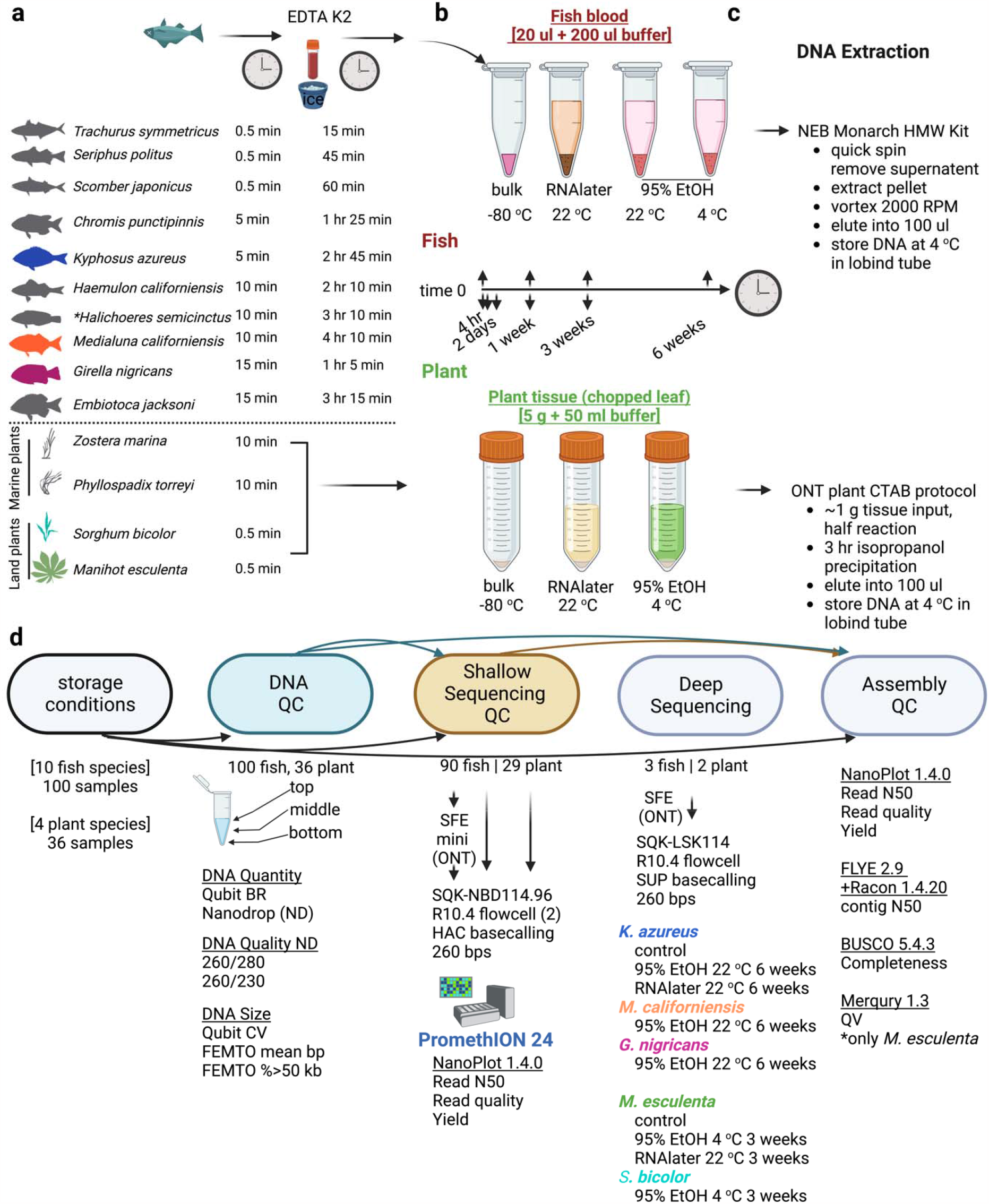
Experimental approach: comparing impacts of storage conditions on DNA QC, sequencing, and assembly metrics. a) Ten fish species and four plant species were collected and b) preserved using various storage conditions for up to 6 weeks (fish) or 3 weeks (plants). c) HMW DNA extraction methods with slight modifications. d) Workflow for assessing impacts of storage conditions on DNA QC and shallow sequencing performance. A subset of species (indicated by color coding) at their storage extremes were fully sequenced and assembled.

Fish were preserved in three distinct storage conditions: 95% EtOH at 4°C, 95% EtOH at 22°C, and RNAlater at 22°C while plants were only preserved under two storage conditions due to constraints on tissue quantity: 95% EtOH at 4°C and RNAlater at 22°C. A total of nine fish and four plant species were included in the analysis of storage conditions on DNA quality metrics, with one fish species (10 samples) excluded due to insufficient blood (Supplementary Table 1). Seven plant samples from later storage time points were excluded from sequencing due to insufficient DNA (Supplementary Table 2). Comprehensive quality control (QC) and sequencing metrics are available in Supplementary Table 2, and statistical analysis across Supplementary Figures 1-4. Source data is available in Supplementary Table 2.

First, we assessed the impact of storage conditions on DNA extraction yield by Qubit and purity by Nanodrop A260/280 and A260/230. All metrics remained stable for fish blood across all storage conditions through six weeks, with yields consistently greater than the liquid nitrogen control (Extended Data Fig 1a-i, Extended Data Fig 2a-c). Greater variability was observed with plant tissue; EtOH 4°C produced consistent yields as a function of time until week three, while RNAlater 22°C produced higher yields at four hours in storage buffer than the liquid nitrogen control, but decreased significantly across the remaining timepoints (P=0.0071, Fs=11.40) (Extended Data Fig 4a,b). DNA A260/230 purity was suboptimal in weeks one and three for plants (Extended Data Fig 4c-f).

DNA fragment sizes measured by Femto Pulse produced two useful metrics: average DNA length and percentage of DNA greater than 50 kilobase pairs (kb). For fish, EtOH 4°C performed ‘best’ with no differences compared to the control to six weeks. EtOH 22°C had the greatest variation among time points for the fragmentation methods assessed, with average length differing from control with P=0.0216, Fs=9.667, and weeks three and six trending lower (Extended Data Fig 1i-l, Extended Data Fig 2e,f). Plants retained DNA size with EtOH 4°C up to three weeks, but reduced dramatically in RNAlater 22°C at later time points (Extended Data Fig 4 i-l). Interestingly, same-day four hour storage in RNAlater 22°C yielded longer DNA than the control, which was not observed for EtOH 4°C (Extended Data Fig 4 j,l).

Following DNA QC, 90 fish samples and 29 plant samples were barcoded and run at low coverage on ONT to determine sequencing quality by read quality and read N50 length. Statistical analysis was performed on the land plants only, though the same trends are qualitatively evident in the larger dataset (Extended Data Fig 4 m-x). Encouragingly, read N50 length for fish did not significantly change with storage time in EtOH 4°C and EtOH 22°C, and was only negatively associated with extended storage times for RNAlater 22°C (Extended Data Fig 1s-u) (Extended Data Fig 2g,j). Reduction in N50 length with storage time was significant in plants for both EtOH 4°C (P=0.0417, Fs=7.600) and RNAlater 22°C (P=0.0417, Fs=7.600) (Extended Data Fig 4 m-p). For read quality, all fish treatments were stable across timepoints, with EtOH 4°C actually showing increased quality by week six (Extended Data Fig 1v-x). Quality scores among the plant samples were stable for EtOH 4°C treatment, but RNAlater 22°C showed reduced quality by day two (Extended Data Fig 3 q-t). DNA fragment sizes did not predict read N50 length for fish, whereas for plants, both DNA yield and fragment size estimates had a positive correlation with read N50 length (Supplementary Table 3). While we observed no association between sequencing yield and N50 length (Extended Data Fig 5a), we found a strong positive correlation between library read quality and read N50 length for both fish and plants (Extended Data Fig 5b).

We were interested in whether size selection could rescue degraded samples, and evaluated the performance of the ONT short fragment eliminator (SFE) kit on the fish dataset of 90 samples. The SFE kit had a positive impact on sequencing N50 (P=0.0443) (Extended Data Fig 6a). The N50 length increased in 63.3% of libraries by an average of 27%. The max increase was 318% (4,092bp to 17,094 bp) and 15.5% of libraries increased N50 length by at least 100% (Extended Data Fig 6b). The application of the SFE kit had the greatest positive impact on highly fragmented libraries, whereas libraries that previously demonstrated a high sequencing N50 length were more negatively impacted by an SFE kit step (Extended Data Fig 6d). Because of the overall positive effect, all DNA samples prepared for deep sequencing were processed with the SFE kit.

For five species, we deep sequenced and assembled high quality reference genomes at the furthest storage time point: three previously unsequenced species of fish (*Medialuna californiensis, Girella nigericans, Kyphosus azureus*) at EtOH 22°C for six weeks, and the two terrestrial plants (*Sorghum bicolor, Manihot esculenta*) in EtOH 4°C for three weeks. Comprehensive sequencing and assembly statistics are presented in Table 1. Notably, all EtOH stored samples assembled with contig N50 lengths greater than 6 Mb. *K. azureus* EtOH 22°C assembled with a 13.82 Mb contig N50 length and 98.6% Benchmarking Universal Single-Copy Orthologs (BUSCO) complete genes (C:98.6%) from 33.9X coverage of the genome, comparable to the liquid nitrogen control with a 21.78 Mb contig N50 length and C:98.8% from 28.4X coverage. *M. esculenta* EtOH 4°C sequence produced a 10.4 Mb N50 with C:99.2% from 58X coverage, while the control yielded 16.5 Mb N50 and C:99.0% from 52.8X coverage. Two included RNAlater 22°C assemblies were reduced in quality: *K. azureus* 5.1Mb N50 length, C:98.7% and *M. esculenta* 0.4 Mb N50 length, C:99.0%. As an additional case study, Illumina reads associated with the same *M. esculenta* cultivar were used to estimate the sequence quality using Merqury statistics to ascertain base-level accuracy among the cassava assemblies^7^. The control, EtOH 4°C, and RNAlater 22°C ONT assemblies were QV= 47.5, 43.8, and 32.1, respectively and showed negligible improvement from Illumina polishing (QV= 49.3, 44.6, and 32.5).

**Table 1.** Sequencing and assembly results of fish and plant samples from extended storage conditions.

| Species | storage |  |  | sequencing |  |  |  |  | Assembly |  |  |  |  |
| --- | --- | --- | --- | --- | --- | --- | --- | --- | --- | --- | --- | --- | --- |
|  | Stored | °C | Weeks | N50_s | N50_d | Long | Q_avg | cov(x) | N50 | contig | C(%) | QV_n | QV_hyb |
| Mcal | ET | 22 | 6 | 3.4 | 9.8 | 176.5 | 18.7 | 58 | 6.5 | 1761 | 98.7 |  |  |
| Gnig | ET | 22 | 6 | 3.5 | 15.2 | 139.4 | 17.9 | 37.8 | 9.7 | 1441 | 98.5 |  |  |
| Kazu | LN | -80 | 0 | 16.9 | 54.4 | 418.9 | 20.1 | 28.4 | 21.8 | 641 | 98.8 |  |  |
| Kazu | ET | 22 | 6 | 15.8 | 20.5 | 183.4 | 19.1 | 33.9 | 13.8 | 792 | 98.6 |  |  |
| Kazu | RN | 22 | 6 | 13.7 | 15.1 | 225.5 | 19 | 11.2 | 5.1 | 795 | 98.7 |  |  |
| Mesc | LN | -80 | 0 | 21.6 | 38.7 | 367.8 | 19.2 | 52.8 | 16.5 | 460 | 99.0 | 47.5 | 49.3 |
| Mesc | ET | 4 | 3 | 13.9 | 24.1 | 296.2 | 17.8 | 58.0 | 10.4 | 638 | 99.2 | 43.8 | 44.6 |
| Mesc | RN | 22 | 3 | 1.9 | 6.2 | 392.3 | 17.2 | 37.3 | 0.4 | 18881 | 99.0 | 32.1 | 32.5 |
| Sbic | ET | 4 | 3 | 9.1 | 18.9 | 394.9 | 19.4 | 54.8 | 8.7 | 845 | 94.4 |  |  |

Species: *M. californiensis* (Mcal), *G. nigericans* (Gnig), *K. azureus* (Kazu), *M. esculenta* (Mesc), *S. bicolor* (Sbic)
Stored: ET (95% EtOH), RN (RNAlater), LN (Liquid Nitrogen control)
N50\_s: sequencing read N50 (kb) from shallow seq approach (Nanoplot) of untreated DNA
N50\_d: sequencing read N50 (kb) from deep seq approach (Nanoplot) of DNA processed through SRT
Long: longest read observed in sequencing (kb)
Q\_avg: mean quality score of library (Nanoplot)
cov(x): estimated sequencing coverage from ONT
N50: the assembly N50 (Mb)
contig: total number of contigs from assembly

Both DNA fragment length and read N50 length from shallow sequencing predicted assembly outcomes from deep sequencing. DNA fragment size is positively correlated with deep sequencing read N50 length (P=0.0101, R²=0.6354) and resulting assembly contig N50 length (P=0.0034, R²=0.7292; linear model) (Extended Data Fig 7a). Shallow sequencing N50 lengths is also positively correlated with deep sequencing read N50 length (P=0.0174, R²=0.5783) and assembly contig N50 length (P=0.0220, R²=0.5507). Of note, shallow sequencing N50 always underestimates deep sequencing N50 which we speculate is due to additional pipetting requirements for barcoding. Nonetheless, their correlation suggests that shallow sequencing can be a powerful QC method for evaluating sequencing suitability of large sample sets where methods like Femto Pulse are time-consuming or unavailable (Extended Data Fig 7b). Genome coverage did not predict assembly N50 length, indicating both that the range of sequencing depths represented in this dataset were more than sufficient. The strongest predictor of assembly length was the deep sequencing N50 length (P=0.0003, R²=0.8649) (Extended Data Fig 7c,d).

Here we show that 95% EtOH is a suitable storage buffer for both fish blood and plant tissue for up to six and three weeks at 22°C and 4°C, respectively, as validated by exceptionally high-quality assemblies (assembly N50 length >1Mb and BUSCO completeness >90%). Furthermore, for a single plant sample stored in 95% EtOH for three weeks at 4°C, we show that ONT sequencing data alone without Illumina polishing can generate a highly accurate consensus reference genome with a QV of 43.8. Overall, we recommend that for animals with nucleated blood, whole blood should be initially stored in an EDTA K2 tube on ice for up to four hours and transferred to 95% EtOH for up to six weeks at 22°C, but improved when refrigerated. RNAlater produces lower quality sequences with extended storage, but may still generate high-quality assemblies for fish up to six weeks as an option in alcohol-restricted countries. Terrestrial plants may be stored in 95% EtOH and kept on ice or refrigerated up to three weeks, or stabilized in RNAlater for same-day transport only. Marine plants have evolved barriers to liquid intrusion from living in an aqueous environment; they may be stored up to one week in EtOH at 4°C and in RNAlater for same-day transport. Considering the great diversity of metabolic content within the plant kingdom, plant tissue storage time should be reduced whenever possible. For ascertaining sequencing suitability within large sample sets, we suggest that QC of DNA is optimally performed by directly sequencing through multiplexing and shallow sequencing on ONT.

## Acknowledgements

We thank the Michael lab for helpful conversations and input into this study. We thank John Tyson’s work on developing aspects of the short read elimination methodology and sharing on twitter. We also thank the San Diego Botanical Gardens and Salk greenhouses for donating plant tissue. This work was funded by a Bill and Melinda Gates Foundation (BMGF) grant (INV-040541) to T.P.M., National Science Foundation (NSF) graduate research fellowship (Fellow #2021321499) to M.L.M. and NSF Postdoctoral Fellowship in Biology “Rules of Life” (Award #2011004) fellowship to J.J.M.

## Author contributions

J.J.M, M.L.M, and T.P.M conceived and designed the experiments. J.J.M carried out the collections, molecular biology, sequencing, and analysis of the fish samples. M.L.M carried out the collection, molecular biology, sequencing, and analysis of the plant samples. J.J.M. and M.L.M. wrote the manuscript. N.A.A. performed the assemblies of fish and plant samples.E.R.M. collected and aggregated data on Femto Pulse. L.T. assisted in molecular biology (extractions). T.P.M supervised the project.

## METHODS

### Sample collection

A total of ten species of marine fish were collected across three different sampling days (September 7th, 9th, and 12th 2022) under IACUC Animal Use Protocol S12219 (Supp Table 1). A total of seven species were collected using a speargun donated by a local fisher. Fish were transported back to shore, euthanized, and blood extracted using a 22 gauge needle and syringe from the caudal vein. The range of transit times from when the fish was first collected until the blood was withdrawn was between 5 and 15 minutes. After blood was drawn, it was dispensed into an EDTA K2 tube and placed on ice. Tubes remained on ice between 1 hr and 5 minutes to 4 hr and 10 minutes for field-collected specimens. An additional three species of fish were collected from an experimental holding tank at Scripps Institution of Oceanography (SIO). These fish were initially collected from the SIO pier and housed in a tank for approximately one month. All three fish had a very short time from when the blood was collected until it was dispensed into the EDTA K2 tube (∼30 seconds or less). These blood tubes also had the shortest time on ice until they were dispensed into the various storage buffers (15 min to 60 min). Fish ranged in size from 12.5 cm to 32 cm in total length and 14.1 g to 519.7 g total mass (Supp Table 1). All fish had sufficient blood for downstream analyses except for *Halichoeres semicinctus*. Ultimately, this fish was excluded from the downstream analyses due to having insufficient blood, but we’ve kept the data here for transparency.

Plants were sourced on the same day (September 12, 2022) from various locations in San Diego, California: Cassava (*Manihot esculenta*) from the San Diego Botanical Gardens (SDBG), Sorghum (*Sorghum bicolor*) from Salk Institute for Biological Studies greenhouse facilities, eelgrass (*Zostera marina*) from Mission Bay, and surfgrass (*Phyllospadix torreyi*) from Windansea Beach (Supp Table 1). The eelgrass samples were collected by hand while snorkeling, with California Fish and Wildlife permit S-210200011-21023-001. Surfgrass was collected as fresh wrack (floating with rhizomes intact, not attached to reef). To achieve relative consistency in tissue age and condition across treatments of the same plant, leaves were cut into 1 cm segments and aggregated before being allocated between storage conditions.

### Sample storage

Using a wide bore pipet tip, approximately 20 μl of fish whole blood kept in EDTA K2 tubes on ice was dispensed into 2 ml Eppendorf tubes containing either 200 μl of 95% EtOH or 200 μl of RNAlater. An additional 20 μl sample was dispensed into an empty 2 ml Eppendorf tube on dry ice that was then immediately stored at -80 °C and used as the control. For samples in buffers, 95% EtOH stored samples were stored at two temperatures of either 4°C (representing refrigeration) or 22°C, representing room temperature. The RNAlater stored samples were only stored at 22°C. Preliminary pilot experiments suggested that ethanol would have a higher performance so we opted to include more temperature treatments with this buffer. We included RNAlater however, because some countries in the world restrict or prohibit all forms of ethanol thus alternative storage buffers are needed in those cases. Samples were then stored for 1 week, 3 weeks, and 6 weeks. At each time point, samples were immediately processed for DNA extraction

At the time of plant tissue sampling, young leaf tissue was added to one of three treatments: liquid nitrogen, 95% ethanol on ice, or RNAlater at ambient temperature. For the ethanol and RNAlater, ∼5g tissue was added to 50mL solvent in nonreactive glassware for storage, in accordance with the suggested RNAater tissue to solvent ratio. Ethanol samples were stored at 4°C and RNAlater samples at room temperature in the dark to reduce oxidation. At each time point, samples were removed from their solvent, blotted with KimWipes and flash frozen in liquid nitrogen for tissue grinding. The same-day time point (time 0.17) was captured 4 hours after collection, once samples reached the lab.

### Sample extraction

Prior to extraction of fish samples stored in buffers, tubes were spun down for 2 minutes at 5000 rpm in a centrifuge. The storage solution supernatant was carefully removed with a pipet. Samples were then processed following the NEB Monarch HMW DNA extraction kit for cells and blood (New England Biolabs, Ipswich, MA, Cat#T3050L).

For buffer stored samples, we followed the ‘fresh nucleated blood’ protocol. For the control samples which had frozen whole blood without buffer, we followed the ‘frozen nucleated blood’ protocol. During the lysis step, we used the highest recommended setting of 2000 RPM. For all fish samples we followed the ‘standard input’ for the various buffer formulations. In the end, we eluted with 100 μl of elution buffer. Final elutions of DNA were allowed to rest at 4°C for at least one week before QC was performed. We deviated from the protocol in that we did not do any sort of pipet mixing as they recommend a 5-10x pipet mix to shear the DNA and enable it to better go into solution. We did this to maximize DNA length.

Plant samples were ground by mortar and pestle in liquid nitrogen to a fine powder and approximately 1g of sample carried forward into extraction, estimated by a half teaspoon scoop. HMW DNA was extracted using the Oxford Nanopore plant extraction protocol for Arabidopsis available to the ONT community, which uses components of the QIAGEN Blood and Cell Culture DNA Midi Kit (QIAGEN Cat#13343). This protocol was adapted for half-reactions in 20 mL lysis buffer to increase throughput by eliminating the combination of two lysed samples per column. Additionally, the final isopropanol precipitation was reduced to 3 hours for all extractions to accommodate samples in RNAlater. The experimental design of collection, storage, and processing is included in Figure 1, which was made using Biorender with a paid license. Statistical analyses and figures made using either R or Prism version 9.4.1 (paid license)

### DNA QC

All DNA was allowed to rest at 4°C for at least one week before quantifying to ensure full solubilization of DNA. Endpoint measures were assessed for DNA QC spanning DNA yield, DNA purity, and DNA fragment size. DNA was quantified using both the Qubit (dsDNA quantitation) and Nanodrop. For the Qubit, 1 μl of DNA from the top, middle, and bottom of each tube was added to the Qubit Broad Range (BR) buffer (Cat#) and quantified. Samples which were below detection were then processed using the Qubit High Sensitivity (HS) kit (Cat#) with 2 μl of DNA. For nanodrop readings, 2 μl of DNA from the top, middle, and bottom of the tube was processed on the spectrophotometer. The concentration in ng/μl, absorbance 260/280, and 260/230 measurements were all recorded. DNA yield was determined by multiplying the total elution volume (100 μl) by the mean DNA concentrations of the top, middle, and bottom of the tube. DNA concentrations were measured with both the Qubit BR kit and Nanodrop, although only the Qubit measurement was used in the final comparison. DNA purity was assessed on the basis of both A260/280 ratios and A260/230 ratios. Again the mean values from the top, middle, and bottom of the tube was used as the endpoint. DNA fragment size can be challenging to measure so we used a three pronged approach. First we used the Coefficient of Variation from the DNA concentration measurements obtained from the Qubit BR kit (top, middle, and bottom of tube). If the DNA is very long, it may not be as homogeneously distributed in solution thus one’s measurements will be more variable. The coefficient of variation (CV) from repeated DNA concentration measures may indicate the homogeneity of DNA in solution with a high CV associated with long fragments. For the other approaches, we directly measured DNA fragment sizes using an automated pulsed-field capillary electrophoresis system (Femto Pulse Cat# M5330AA, Agilent, Santa Clara, CA) which can reliably measure DNA fragments up to 165 kb in length. We curated two endpoint measures with the Femto pulse; first the mean read length which is automatically generated in the program and second the percent of DNA greater than 50 kb. For the later measurement, we used the ‘smear analysis’ function built into the Femto pulse analysis program to quantify DNA measurements between 1-10 kb, 10-25 kb, 25-50 kb, 50-100 kb, and greater than 100 kb. These values can be found in the metadata in Supplement. To simplify the analyses, we used a final measure of the % of DNA between 50-100 kb and greater than 100 kb. Previous sequencing experience in the lab has shown that sequencing N50s greater than 20-30 kb is usually sufficient to obtain high quality reference genomes during assembly.

### Sample exclusion criteria

Whole blood extracted from ten species of fish was stored at different conditions (95% EtOH 4°C, 95% EtOH 22°C, and RNAlater 22 °C) for 0, 1, 3, and 6 weeks (Figure 1a). Across all 100 samples, 5 samples from one species (*H. semicinctus*) had insufficient DNA quantity by Nanodrop (<10 ng/ul), thus all 10 samples from this species were excluded from analysis and sequencing. This fish had insufficient blood at the start and required addition of PBS to dilute the sample. Leaf tissue from 4 species of plants was stored in 95% EtOH at 4°C or RNAlater 22°C for 0, 0.17(4hr), 2, 7, and 21 days. Of 36 samples, 5 had insufficient DNA (*Z. marina* RNAlater22°C 2 days, 1 week, 3 week; *P. torreyi* RNAlater 1 week and 3 week) while another 2 samples (*Z. marina* EtOH4°C 1 week and 3 weeks) had poor quality DNA thus were excluded from further analysis (Figure 1a).

### Shallow sequencing

To assess the influence of sample storage conditions on DNA sequencing, sequencing libraries were made using the newest barcode ligation chemistry (Kit 14) from Oxford Nanopore Technologies (ONT). Samples were processed together to increase throughput and minimize variability during library preparation from pipet shearing of DNA etc. Sequencing libraries were made using either 400 ng (plant) or 800 ng (fish) of DNA as input according to the ONT protocol. The Native Barcoding Kit 96 V14 SQK-NBD114.96 was used to construct libraries either 90 (fish) or 29 (plant) at a time. Each unique pool was sequenced separately on a new Promethion 10.4.1 flow cell. MinKnow version 22.1 was used to process and samples were basecalled using the high accuracy basecalling. At termination of sequencing, libraries were then processed using the NanoPlot tool to generate metrics for each sample including average read length, N50, total reads, total bases, and average read quality.^8^

### Evaluation of sample storage on DNA QC and sequencing

Our goals were to determine if sample storage had an impact on DNA quality and if sample storage had an impact on sequencing results. A third goal was to also determine which DNA QC metrics are useful for predicting sequencing success. To determine which storage condition (95% EtOH 22°C, 95% EtOH 4°C, and RNAlater 22°C) performed best for fish samples, we specifically performed a one way Friedman test with repeated measures for the four time points (0 week, 1 week, 3 weeks, and 6 weeks) followed by a multiple comparisons test against the control (week 0, dry ice, -80°C stored) ^9^. Our design had 4 groups and 9 replicates (species) for the fish samples. For plant samples, we compared 95% EtOH 4°C, and RNAlater 22°C across a total of 5 time points (0, 4 hr, 2 days, 7 days, and 3 weeks) and we had 4 replicates (species). We had to use a non-parametric test because sample groups failed to pass normality testing with Shapiro-Wilk ^10^. Multiple comparison correction was performed using Two-stage step-up method of Benjamini, Krieger and Yekutieli with an alpha of 0.05.^11^ Each of the three storage conditions were independently analyzed to determine if sampling time had an impact. The statistical results could then be compared across the three storage buffer and temperature combinations.

### Impact of short read exclusion kits on sequencing performance

All 90 fish samples were sequenced with and without a size selection (ONT SFE) (Cat # EXP-SFE001). Specifically, approximately 5 ug of DNA was used as input in a total of 25 ul which was then matched with 25 ul of SFE buffer. This is a slight modification (miniaturization of the protocol which recommends starting with at least 50 ul) and the sequencing read N50 values compared using the Wilcoxon matched-pairs signed rank test (two-tailed) as they did not pass the Shapiro-Wilk normality test (P=0.0005 and P=0.0012). The percent increase in N50 was calculated for the 90 pairs by taking (N50 SFE - N50 noSFE)/(N50 noSFE) * 100. A positive value indicates an increased N50 as a result of using SFE.

### Evaluation of DNA QC on sequencing QC

Fish and plant samples chosen for higher coverage whole genome sequencing were additionally size selected using ONT size selection (Oxford Nanopore Technologies, Cat# EXP-SFE001) to deplete short fragments under 25 kb. We performed deep sequencing on a subset of plant and fish samples to evaluate assembly metrics. We did deep sequencing on three species of fish including *K. azureus, M. californiensis*, and *G. nigricans*. This included samples stored at 95% EtOH at 22C for 6 weeks. For *K azureus*, we also sequenced the control and the sample stored in RNAlater at 22°C for 6 weeks. For plants, we sequenced the samples stored in 95% EtOH at 4°C for 3 weeks from two important food crops, *M. esculenta* (cassava) and *S. bicolor* (sorghum). For cassava, we also sequenced the control and the samples stored in RNAlater at 22°C for 3 weeks (Table 1). For assembly validation, we followed the Vertebrate Genome Project (VGP) guidelines.^4^ Data quality control was performed with FastQC version 0.11.0 and NanoPlot 1.40.0.^8^ Unassembled genome analysis and size estimation was completed using GenomeScope 2.0.^12^ Samples were assembled using Flye version 2.9.^13^ All nanopore only assemblies were polished with the raw nanopore reads using Racon version 1.4.20.^14^ In an separate assembly, the cassava sample was additionally polished with Pilon version 1.24 after the initial assembly.^15^ All polishing utilized Minimap2 version 2.21.^16^ Final assembly metrics were calculated using assembly-stats version 1.0.1 and completeness was estimated using BUSCO version 5.4.3.^17^ The BUSCO databases eudicots_odb10, liliopsida_odb10, and actinopterygii_odb10 were used for cassava, sorghum, and fish samples, respectively. QV was calculated using Meryl version 1.3 and Merqury version 1.3.^7^

## Notes

### Competing Interest Statement

The authors have declared no competing interest.

