## Supplemental figures for "Generating high quality reference genomes from field collected specimens by optimizing preservation"

### Extended Data Figures

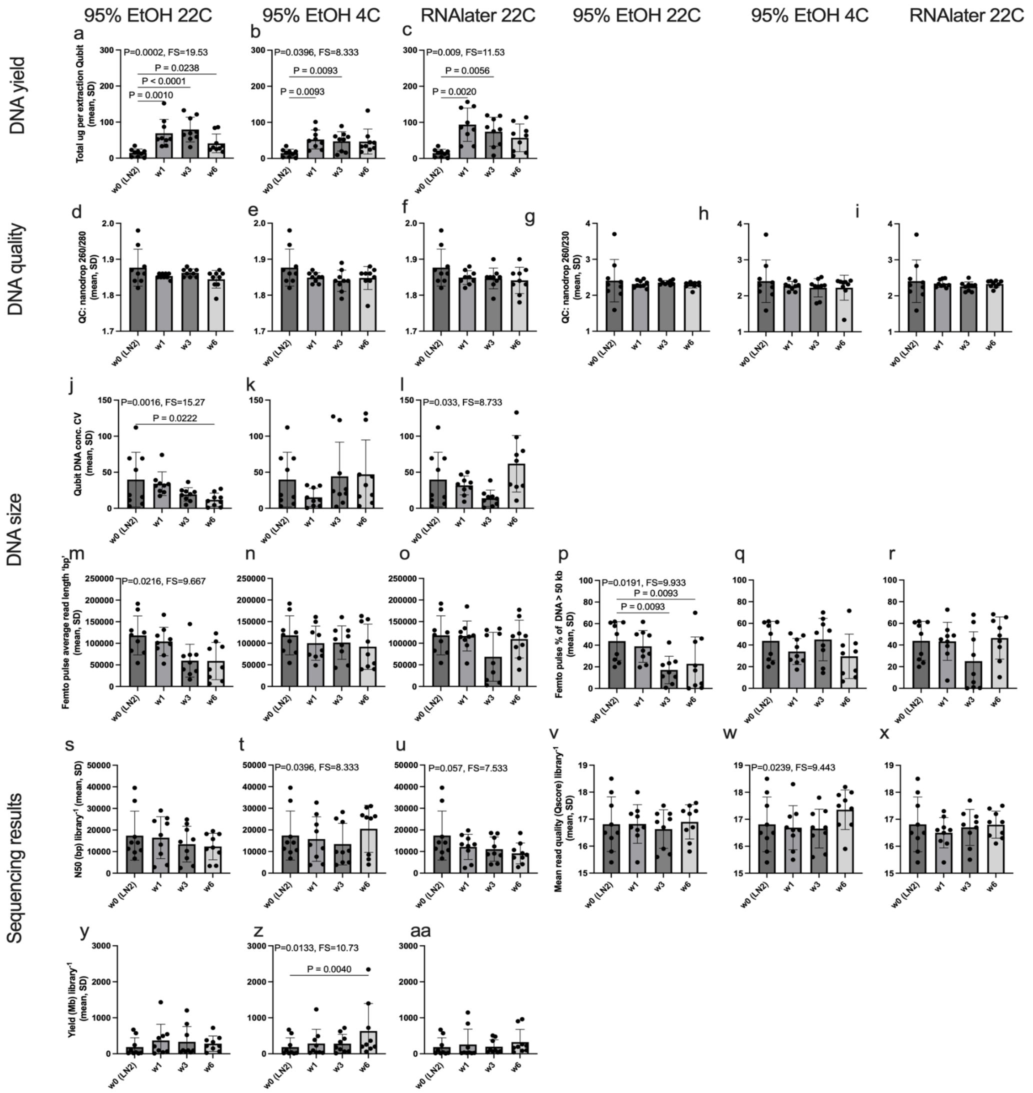


Extended Data Figure 1. Impacts of sample storage buffer, temperature, and time on fish DNA quality and sequencing. DNA samples from 9 fish (1 fish excluded) were sequenced on Oxford Nanopore Promethion LSK114. Variation of DNA yield (total ug) as measured by Qubit BR kit amongst samples stored in a) 95% EtOH 22C, b) 95% EtOH 4C, and c) RNAlater 22C. Variation in DNA quality 260/280 as measured by Nanodrop amongst samples stored in d) 95% EtOH 22C, e) 95% EtOH 4C, and f) RNAlater 22C. Variation in DNA quality 260/230 as measured by Nanodrop amongst samples stored in g) 95% EtOH 22C, h) 95% EtOH 4C, and i) RNAlater 22C. Variation in DNA fragment size as measured by the coefficient of variation of Qubit readings from top, middle, and bottom of tube across j) 95% EtOH 22C, k) 95% EtOH 4C, and l) RNAlater 22C. Variation in mean DNA fragment size (bp) as measured by femto pulse across m) 95% EtOH 22C, n) 95% EtOH 4C, and o) RNAlater 22C. Variation in DNA fragment sizes greater than 50kb as measured by femto pulse across p) 95% EtOH 22C, q) 95% EtOH 4C, and r) RNAlater 22C. Variation in sequencing results as measured by NanoPlot output for N50 across s) 95% EtOH 22C, t) 95% EtOH 4C, and u) RNAlater 22C; mean read quality across v) 95% EtOH 22C, w) 95% EtOH 4C, and x) RNAlater 22C; and total yield across y) 95% EtOH 22C, z) 95% EtOH 4C, and aa) RNAlater 22C

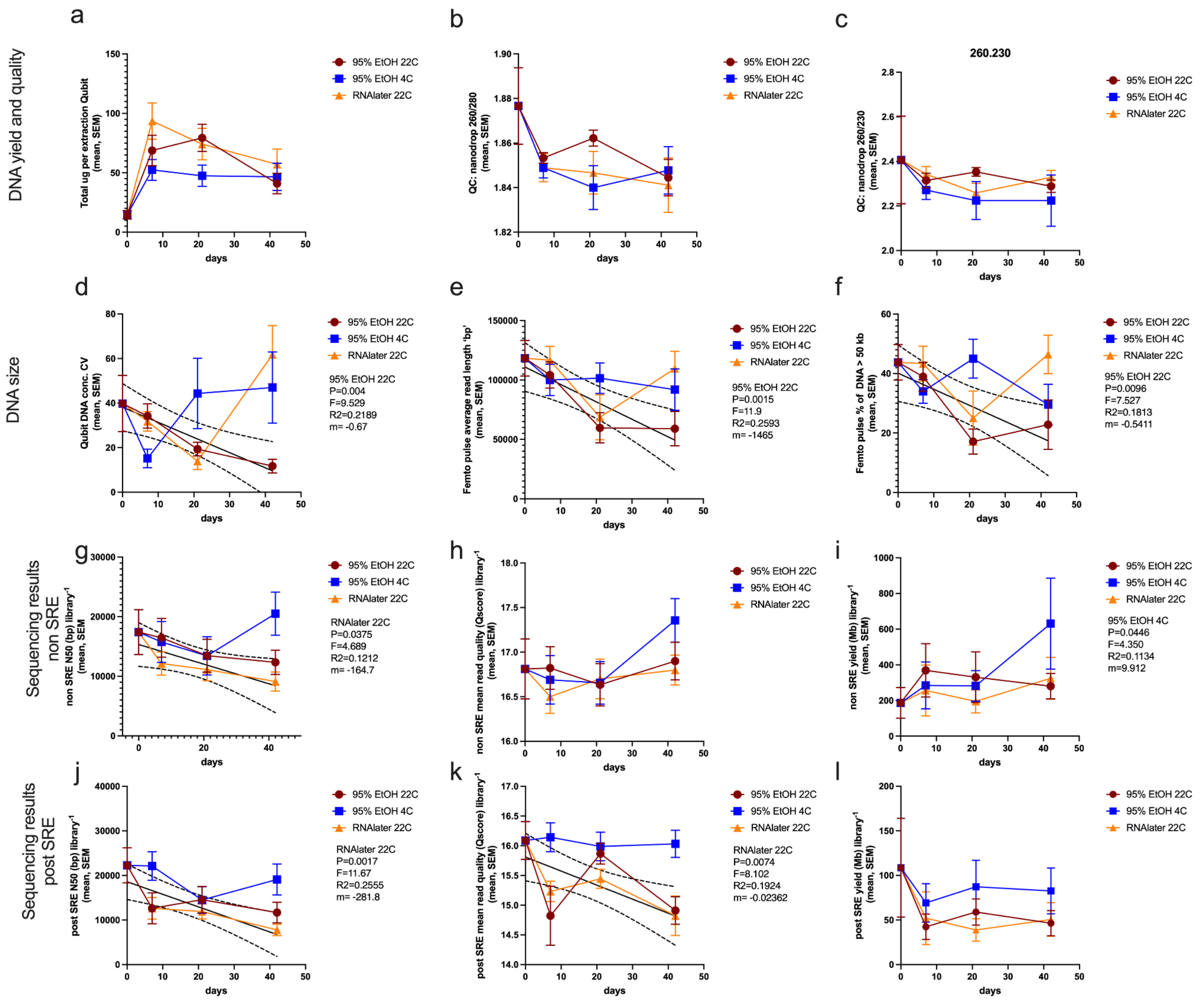


Extended Data Figure 2. Impact of storage time on DNA QC metrics and shallow sequencing results. Linear regression analysis performed on 9 fish species as replicates across three storage conditions (95% EtOH 22C, 95% EtOH 4C, and RNAlater 22C). a) DNA yield as measured by total ug per reaction, DNA quality as measured by b) nanodrop 260/280 ratio and c) nanodrop 260/230 ratio. DNA size was measured using d) the CV from Qubit measurements of the top, middle, and bottom of the tube, e) FEMTO pulse average bp, and f) FEMTO pulse percent of DNA larger than 50 kb. Shallow sequencing performed on original DNA and DNA processed with the ONT short read eliminator kit. Sequencing metrics for original DNA was obtained for g) read N50, h) average read quality score, and i) total yield in Mb. Sequencing metrics for DNA processed using the SRE kit was obtained for j) read N50, k) average read quality score, and l) total yield in Mb.

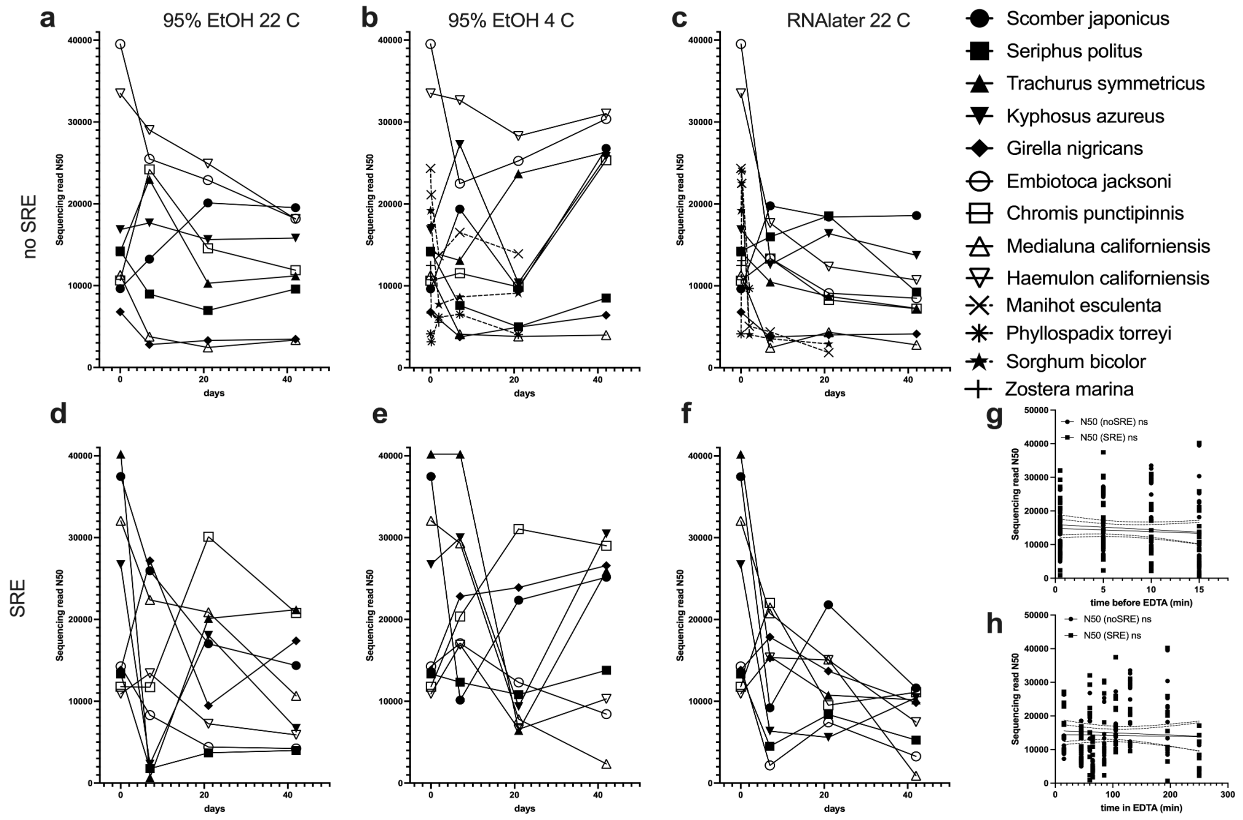


Extended Data Figure 3. Variation of sequencing N50 across samples as a result of storage buffer, time, and use of short read eliminator. Sequencing N50 results of untreated DNA from samples stored in a) 95% EtOH 22 oC (9 fish species), b) 95% EtOH 4 oC (9 fish and 4 plant species), and c) RNAlater 22 oC (9 fish and 4 plant species). Plants are indicated by a dashed line while fish have solid line. Sequencing N50 results of DNA processed through the ONT SRE kit d) 95% EtOH 22 oC (9 fish species), e) 95% EtOH 4 oC (9 fish species), and f) RNAlater 22 oC (9 fish species). Impacts of blood collection methods also compared to sequencing N50. Specifically, g) impact of time that blood remained inside fish prior to being dispensed into an EDTA tube on sequencing N50 (30 seconds to 15 minutes) and h) impact of time in which EDTA blood tube was incubated on ice prior to being aliquoted to storage buffers (15 minutes to 4 hr 10 min). (g-h) Statistical significance tested with linear model and Spearman correlation.

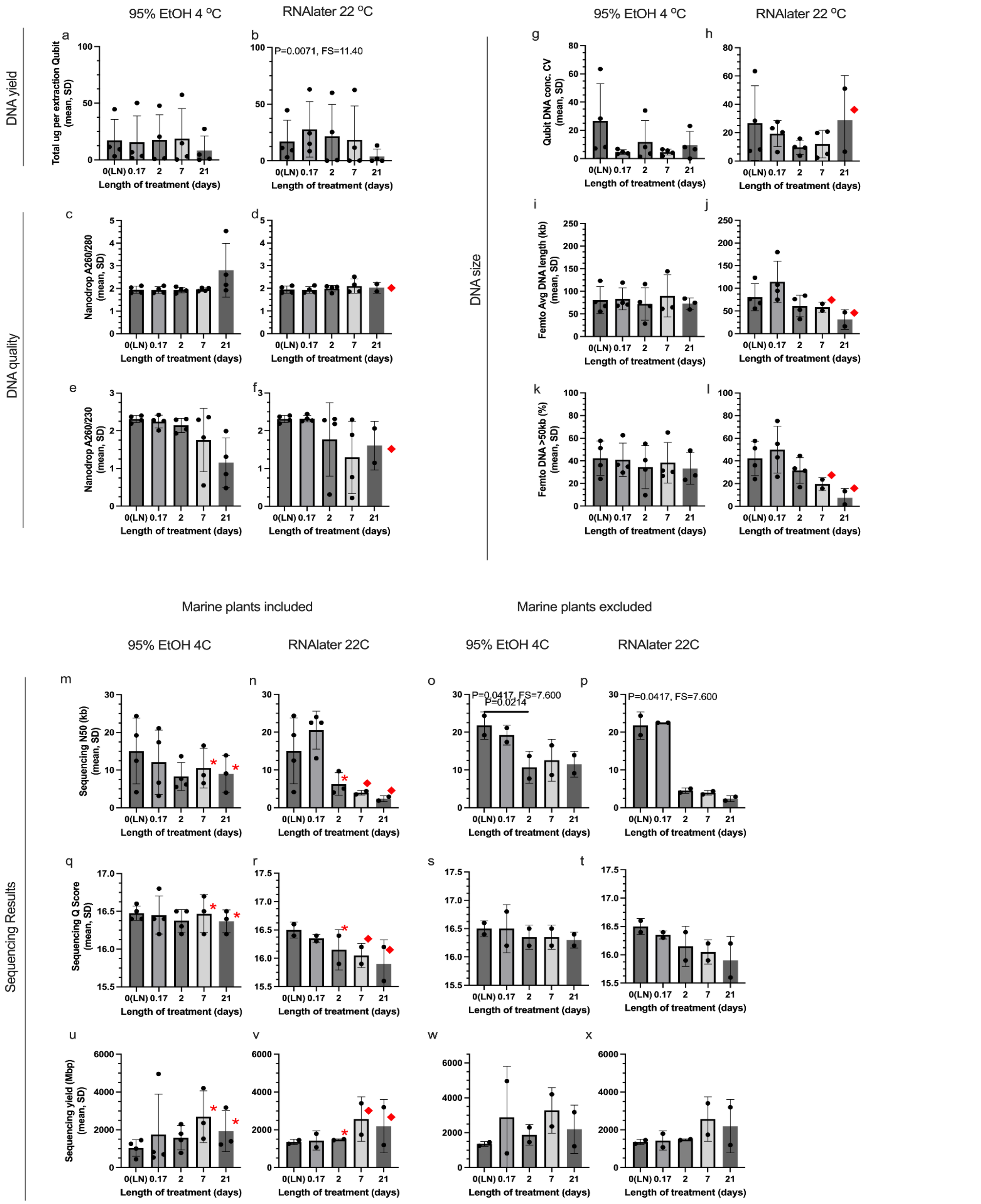


Extended Data Figure 4. Impacts of sample storage buffer, temperature, and time on plant DNA quality and sequencing. DNA samples from 4 plants were sequenced on Oxford Nanopore Promethion LSK114. Given the low yields for some later marine plant (*Z. marina* and *P. torreyi*) time points, the sequencing statistics are presented with and without these plants present, and statistical analysis run on the terrestrial plants only.

Variation of DNA yield (total ug) as measured by Qubit BR kit amongst samples stored in a) 95%ETOH at 4C and b) RNAlater at 22C. Variation in DNA quality 260/280 as measured by Nanodrop amongst samples stored in c) 95%ETOH at 4C and d) RNAlater at 22C. Variation in DNA quality 260/230 as measured by Nanodrop amongst samples stored in e) 95%ETOH at 4C and f) RNAlater at 22C. Variation in DNA fragment size as measured by the coefficient of variation of Qubit readings from top, middle, and bottom of tube across g) 95%ETOH at 4C and h) RNAlater at 22C. Variation in mean DNA fragment size (bp) as measured by femto pulse across i) 95%ETOH at 4C and j) RNAlater at 22C. Variation in DNA fragment sizes greater than 50kb as measured by femto pulse across k) 95%ETOH at 4C and l) RNAlater at 22C. Variation in sequencing results as measured by NanoPlot output for N50 across all sequenced samples for m) 95%ETOH at 4C and n) RNAlater at 22C, and across only terrestrial samples for o) 95%ETOH at 4C and p) RNAlater at 22C. NanoPlot mean read quality across all sequenced samples for q) 95%ETOH at 4C and r) RNAlater at 22C, and across only terrestrial samples for s) 95%ETOH at 4C and t) RNAlater at 22C. NanoPlot total yield across all sequenced samples for u) 95%ETOH at 4C and v) RNAlater at 22C, and across only terrestrial samples for w) 95%ETOH at 4C and x) RNAlater at 22C.
* indicates the time point contains a single marine plant sample below QC detection limit or omitted from sequencing, and the time point is omitted from the statistical analysis in that graph

♦ indicates the time point contains two marine plant samples below QC detection limit or omitted from sequencing, and the time point is omitted from the statistical analysis in that graph

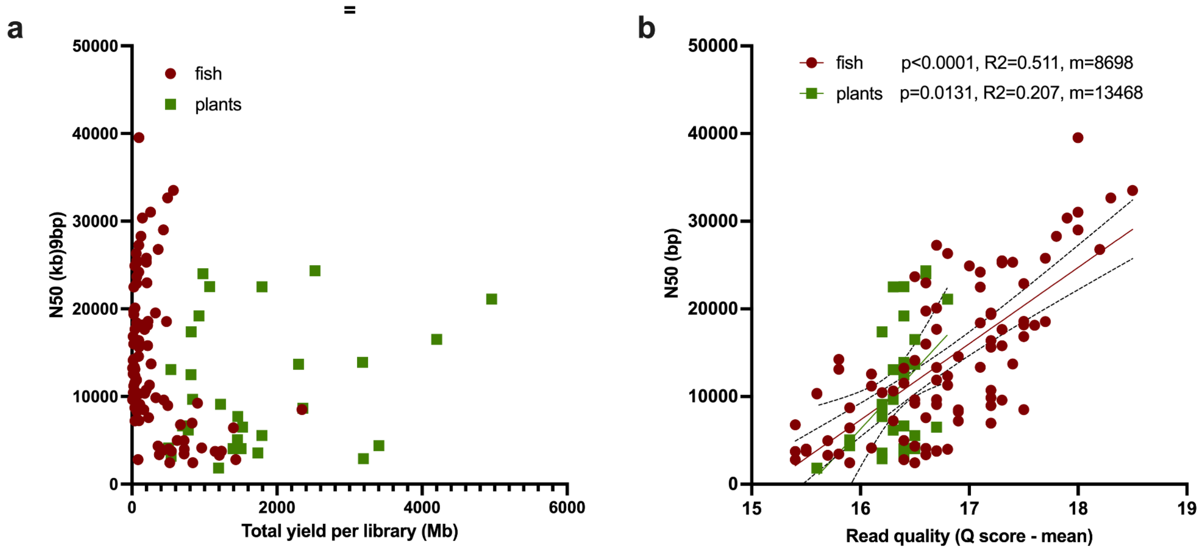


Extended Data Figure 5. Impacts of sequencing metrics such as a) total library yield and b) mean read quality per library on N50 scores. All DNA from 90 fish samples and 29 plant samples were run on two separate sequencing runs (R000-469, and R000-471, respectively) using the LSK114 native barcoding kit. Average read quality, total yield, and N50 scores were generated from NanoPlot. Linear regression with 95% CI shown in panel b.

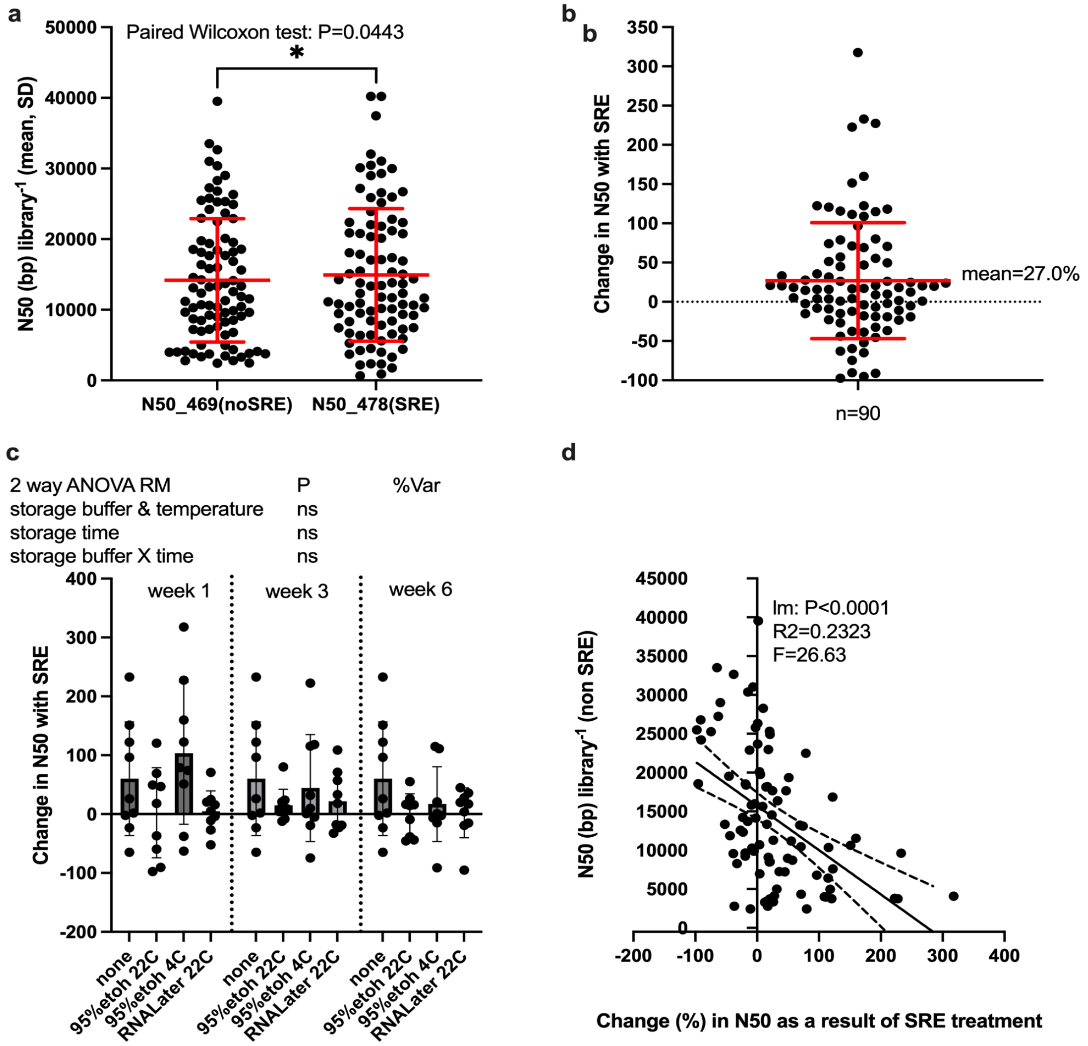


Extended Data Figure 6. Impacts of short read exclusion kit on sequencing performance as measured by sequencing read N50 generated by NanoPlot. a) All 90 DNA fish samples processed in two sequencing runs with and without SRE with pairwise comparison of N50 (Paired Wilcoxon test; P<0.05 *). b) Percent change of N50 as a result of using the SRE kit (positive indicates that the N50 increased with using SRE), c) comparison of sample storage groups on benefit of SRE, and d) impacts of change in N50 as a result of using SRE on the original N50 value (preSRE). Linear model for assessment.

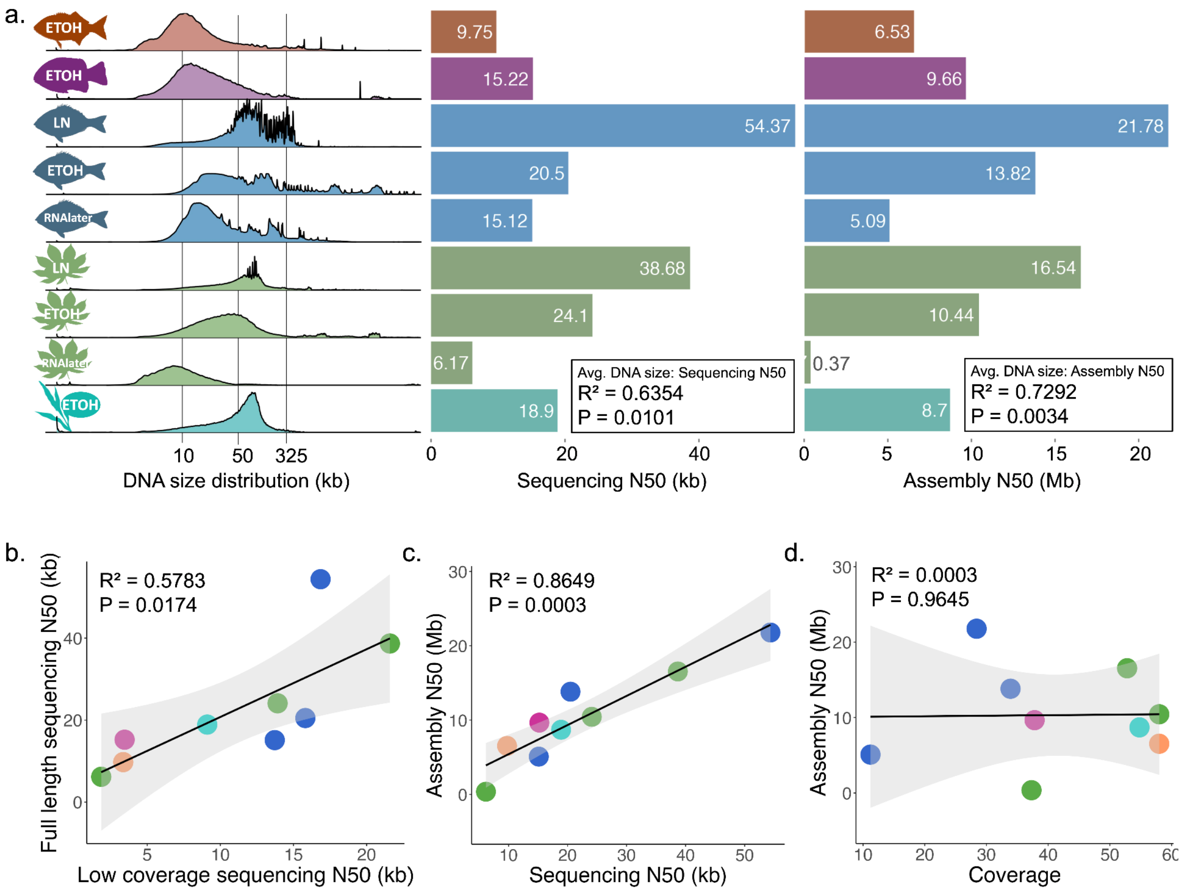


Extended Data Figure 7. Sequencing and assembly N50 statistics for fully sequenced samples demonstrate successful assembly at furthest time points for 3 fish species (*M. californiensis, G. nigiricans, K. azureus*) at 6 weeks in 95% ETOH at 22C and 2 plant species (*M. esculenta, S. bicolor*) at 3 weeks in 95% ETOH at 4C. Additionally, *K. azureus* and *M. esculenta* are sequenced from tissue stored in liquid nitrogen (control) and RNAlater at room temperature for 6 weeks and 3 weeks for fish and plants respectively. a) Femto Pulse DNA fragment size distributions are shown alongside the sequencing read length N50 metric generated by Nanoplot and resulting assembly contig N50s from the associated FlyE assembly. b) Correlation between the multiplexed shallow sequencing N50 results and full sequencing N50 results. c) Correlation between sequencing N50s and assembly contig N50 from FlyE assemblies. d) Correlation between genome coverage from passed ONT reads and assembly contig N50 from FlyE assemblies.
